# Methanotrophic Community Detected by DNA-SIP at Bertioga’s Mangrove Area, Southeast Brazil

**DOI:** 10.1101/2020.08.20.255125

**Authors:** Débora do Carmo Linhares, Flávia Talarico Saia, Rubens Tadeu Delgado Duarte, Cristina Rossi Nakayama, Itamar Soares de Melo, Vivian Helena Pellizari

## Abstract

Methanotrophic bacteria can use methane as sole carbon and energy source. Its importance in the environment is related to the mitigation of methane emissions from soil and water to the atmosphere. Brazilian mangroves are highly productive, have potential to methane production, and it is inferred that methanotrophic community is of great importance for this ecosystem. The scope of this study was to investigate the functional and taxonomic diversity of methanotrophic bacteria present in the anthropogenic impacted sediments from Bertioga’s mangrove (SP, Brazil). Sediment sample was cultivated with methane and the microbiota actively involved in methane oxidation was identified by DNA-based stable isotope probing (DNA-SIP) using methane as a labeled substrate. After 4 days of incubation and consumption of 0.7 mmol of methane, the most active microorganisms were related to methanotrophs Methylomonas and Methylobacter as well as to methylotrophic Methylotenera, indicating a possible association of these bacterial groups within a methane derived food chain in the Bertioga mangrove. The abundance of genera Methylomonas, able to couple methane oxidation to nitrate reduction, may indicate that under low dissolved oxygen tensions some aerobic methanotrophs could shift to intraerobic methane oxidation to avoid oxygen starvation.

## Introduction

Mangroves are tropical and subtropical ecosystems in the transition between land and marine environments. Subjected to tidal periodical flooding, mangroves present variable salinity, are muddy, oxygen poor and rich in nutrients and organic matter [1–3]. The environmental variability in mangroves sustains a highly active microbiota, which plays a central role in biogeochemical cycles, soil structure generation and decomposition, and influences the primary production and plant community dynamics [1, 2].

Even though coastal marsh ecosystems are considered net sinks for carbon sequestration, spatial and temporal gradients promote a wide range of biogeochemical and anaerobic conditions, making them important sources of greenhouse gases to the atmosphere [4]. The prevalence of anaerobic conditions partnered with the high organic matter content favors methanogenesis by archaea. Methane generation is in part regulated by competition with sulfate reducing bacteria in periods of higher sulfate concentration, but methanotrophic bacteria and archaea have also a central role in reducing the total net flux of methane to the atmosphere as they are responsible for the oxidation of a significant amount of the gas produced in mangroves [5, 6]. Although anaerobic methanotrophy occurs in mangroves, coupled to sulfate, nitrate or nitrite reduction [7], the consumption of methane by aerobic bacteria is also an important process taking place in the thin oxic layer at the sediment – water interface, submerged leaf sheaths, or associated to plant roots and rhizosphere [1, 8]. Methanotrophic bacteria may also benefit from the interaction with fungi, who produce hydrophobic proteins to reduce surface tension on the hyphae, facilitating the access to hydrophobic gases such as methane [9].

Aerobic methanotrophic bacteria form a group physiologically unique and distinctive for its ability to use methane as a sole source of carbon and energy [10]. Besides their importance in the global cycle of methane and nitrogen, methanotrophic bacteria has potential application in several biotechnological processes, such as remediation of chlorinated solvents, production of polyhidroxyalcanoates (PHA), denitrification, and demethylation of methyl mercury [11], even in adverse oxygen concentrations [12].

Methanotrophic bacteria were originally grouped into Type I, II and X according to phylogeny, cellular ultrastructure, metabolic pathways, and ability to fix nitrogen [5]. As knowledge in aerobic methanotrophic diversity advanced, grouping criteria were revised. In current classification, according to phylogeny and carbon fixation pathways, methanotrophic bacteria are divided into Type I (gamma-proteobacteria using the ribulose monophosphate pathway), Type II (alpha-proteobacteria fixing carbon through the serine pathway) and Type III (*Methylacidiphilae*, in Phylum Verrucomicrobia, using the Calvin cycle to fix carbon derived of methane oxidation) groups. Type I methanotrophs were further divided into Types Ia *(Methylococcaceae*), Ib (*Methylococcaceae*, former Type X), Ic (*Methylotehrmaceae*) and Id (uncultured groups, based on *pmo*A sequences). Type II methanotrophs were also divided into Type IIa (*Methylocystaceae*) and Type IIb (*Beijerinckiaceae*) subgroups [13, 14].

Despite the importance of methanotrophic processes in regulating methane fluxes from coastal marsh ecosystems, studies of methanotrophic diversity in these environments are still scarce compared to research in freshwater and upland soils ecosystems. Brazilian mangroves, considered very vulnerable to damage, correspond to 7 to 8.5% of global mangrove areas and it is discontinuously distributed along the Brazilian coast [15]. Previous studies with Brazilian mangrove samples confirmed the presence of methanogenic archaea and metagenomic analysis showed sulfur metabolism prevalent in microbiomes of polluted and pristine sites [2, 16, 17]. Mesocosm experiments detected changes in bacterial communities induced by oil contamination in mangrove sediments from São Paulo State [18] and reported a preferential enrichment of the aerobic methanotroph Methylococcaeae sequences in the rhizosphere of Rhizophora mangle from Guanabara Bay mangrove, in Rio de Janeiro State [1]. Here we investigate the functional and taxonomic diversity of active methanotrophic bacteria present in oil polluted mangrove sediment samples from Bertioga (São Paulo State, Brazil), through DNA-SIP followed by the construction of 16S rRNA gene libraries, which allows studying the role of active cultured and uncultured bacteria in the oxidation of CH_4._

## Material and Methods

### Sample collection and processing

Surface sediment samples (up to 5 cm below sediment-water interface) were collected from a mangrove located in Bertioga, São Paulo State, in the southeast region of Brazil, in an area chronically contaminated by oil spills [2]. Sediments were sampled at 23°53’49’’S, 46°12’28’’W, from 5 points distanced by 2 to 4 m. Sediment samples, approximately 500 g, were collected with a sterile stainless steel spatula to the depth, sealed in sterile plastic bags and transported in a cool box at 4°C. In the laboratory, samples were homogenized and stored at 4°C for stable isotope probing experiment. Aliquots of 0.25 g of homogenized sediment were immediately stored at −20°C for molecular analyses. At the moment of sampling, values of water salinity, sediment salinity, pH, temperature, conductivity, and dissolved oxygen were 1.74%, 0.48%, 6.58, 22.6°C, 8.84 µs.cm^-1^ and 0.0 mg.L^-1^, respectively.

### Stable isotope probing microcosms

Five grams of homogenized sediment samples (wet weight) were incubated in 100 mL glass bottles filled with 40 mL of NMS medium (ATCC 1306), with salinity of 1.13% adjusted with synthetic reconstituted sea water (S9883, Merck, Germany), and sealed with butyl rubber stoppers and aluminum crimp caps. ^13^C-methane (99% ^13^C, Cambridge Isotope Laboratories, Andover, USA) or ^12^C-methane (Linde, São Paulo, Brazil) was added to a final methane concentration on headspace of 8% (v/v) under sterile conditions, using sterile 0.2 µm hydrophobic PTFE syringe filters. Controls, without methane, were also included. Replicate bottles amended with ^13^C-methane or ^12^C-methane were incubated at 28°C in the dark at 150 rpm. Methane (99% purity) was supplied to the microcosms whenever detected consumption was greater than 95%, up to 8 additions. Before each methane addition, the bottles were flushed with sterile air to reestablish atmospheric conditions. A pair of bottles (^13^C-methane and ^12^C-methane) was subsequently taken for nucleic acid extraction at days 2, 4 and 7, corresponding to methane consumption of 0.2, 0.7 and 1.4 mmol, respectively. Sediment slurry was centrifuged (12.000 x *g*, 40 min, 4°C) and cells in the pellet were immediately stored at −200C.

### DNA extraction and isopycnic centrifugation and fractionation

DNA from the sediment as well as from SIP microcosms were extracted with Power Soil DNA Isolation Kit (Mo-Bio Laboratories Inc, USA) as described in the manufacturer’s protocol. The integrity of DNA was checked on gel electrophoresis and quantified using a NanoDrop ND1000 Spectrophotometer (Thermo Scientific, USA).

Equilibrium (isopycnic) density gradient centrifugation and fractionation were adapted for DNA-SIP from methods for RNA-SIP [19] using cesium trifluoroacetate (CsTFA) gradients, without addition of formamide and with a starting buoyant density (BD) of 1.61 g.mL^-1^. Solutions were prepared by mixing 2.0 g.mL^-1^ CsTFA stock solution (Amersham Biosciences) and gradient buffer described in [20]. Gradients were loaded with 10 μL of DNA (500 ng) and then subject to ultracentrifugation at 64,000 rpm and 20°C for 40 h using the same tubes, rotor and ultracentrifuge described previously [19]. Gradients were fractionated into 100 µL fractions as described previously [19]. Twenty fractions were obtained through each fractionation procedure, which were numbered from 1 (heavier) to 20 (lighter). Buoyant density (BD) of fractions was determined indirectly by measuring refraction index with an AR200 digital refractometer (Reichert Inc., Depew, NY, USA) of each fraction from blank gradients run in parallel containing water instead of DNA. Sample DNA was precipitated overnight from fractions with 500 µL cold isopropanol at −20 °C, followed by centrifugation (14,000 rpm, 30 min, 4°C). Precipitates were washed in 70% cold ethanol (0.5 mL) and re-eluted in 30 µL elution buffer (10 mM Tris-HCl, 1 mM EDTA, pH 8). Total DNA was determined using the PicoGreen^®^ ds-DNA Quantitation Kit (Invitrogen), according to the manufacturer’s instructions.

### Denaturing Gradient Gel Electrophoresis

Phylogenetic diversity of the bacterial communities from the sediment and from each fraction of SIP incubation with ^12^CH_4_ and ^13^CH_4_ at the 4^th^ day was analysed by denaturing gradient gel electrophoresis (DGGE). Bacterial 16S rRNA gene fragments were amplified in a PCR from 1 µL extracted DNA with primers GC338F and 518R [21]. PCR program in a Mastercycler Personal-system (Eppendorf, USA) was 94°C for 5 min, 30 cycles 94 °C for 1 min, 55 °C for 1 min, 72°C for 2 min and a final elongation period of 7 minutes at 72°C. The amplified product (7 μL) was analysed on an 8% polyacrylamide gel with a 45% - 65% denaturing gradient (where the 100% denaturant contained 7 M urea and 40% formamide) that was run for 15 hours at 60°C and 100 V in the Ingeny PhorU2 apparatus (Ingeny International, Goes, The Netherlands). Gel was silver nitrate stained [22] and DGGE profiles were visualized under white light. Comparisons of DGGE profiles were performed by cluster analysis of the banding patterns using Bionumerics™ software (Applied Maths, NV). Dendrograms were constructed by the unweighted pair group method with arithmetic mean (UPGMA) groupings with a similarity matrix based on the Pearson coefficient.

### Clone library and phylogenetic analysis

In order to identify the dominant bacterial species involved in methane oxidation process, 16S rRNA clone libraries were constructed from total DNA of the sediment (GenBank accession No. MT644161-MT644186) and from a heavy and a light DNA fractions from ^13^CH_4_ flask (fractions 12 and 15, respectively) obtained after isopycnic centrifugation (GenBank accession No. MT603661-MT603717). Clone library of *pmo*A genes of the sediment was also done (GenBank accession No. MT596824-MT596880).

Bacterial 16S rRNA gene fragments were PCR amplified in triplicate from 1 µL of total DNA (50 ng) with primers 27F [23] and 1401R [24]. The temperature program was 94°C for 5 min, 30 cycles 94°C for 30 seconds, 55°C for 30 seconds, 72°C for 90 seconds and a final elongation time of 7 minutes at 72°C. Fragments of *pmoA* genes were PCR amplified in triplicate from 1 µL of total DNA (50 ng) with forward primer A189 [25] and reverse primer MB661 [26]. The temperature program was 94°C for 5 min, 30 cycles 94°C for 30 seconds, 55°C for 30 seconds, 72°C for 90 seconds and a final elongation time of 7 minutes at 72°C. Amplified 16S rRNA and *pmo*A gene fragments were purified with Pure Link PCR Purification Kit (Invitrogen), cloned into pGEM-T-Easy (Promega - Madison, Wisconsin, USA) according to the manufacturer’s protocol and transformed into *E. coli* JM109 by heat shock (0°/42 °C for 45 seconds). Cloned inserts were amplified with primers M13F and 1401R for 16S rRNA clones, and with primers M13F and M13R for *pmoA* clones. The temperature program for both reactions was 97°C for 3 min, 40 cycles 94°C for 30 seconds, 60°C for 30 seconds, 72°C for 90 seconds and a final elongation time of 5 minutes at 72°C. The amplified products were purified with Pure Link PCR Purification Kit (Invitrogen) and sequenced (MegaBACE 1000 System) with T7 primer.

Initially, all 16S rRNA sequences were checked for chimeras on the software Bellerophon [25]. Sequences considered putative chimeras or shorter than 540 bp were excluded from further analysis. The 16S rRNA clones were aligned on Mothur v.1.42.0 [26] using the SILVA 138 reference database [27] and the alignment was checked and manually edited for position corrections using the software ARB [28]. A phylogenetic tree was constructed on ARB by the maximum likelihood method with a 1,000 bootstrap analysis. Representative reference sequences of the most closely related members were obtained from the Genbank [29] and Ribosomal Database Project – RDP [30].

A similar approach was used for the analysis of *pmo*A clones. The sequence alignment was performed with the Clustal W [31] within the BioEdit v7.0.9.0 package [32] and the phylogenetic tree was constructed on MEGA 7 [33] using the neighbor-joining method and a 2,000 bootstrap value.

### Chemical analysis

Methane was measured by headspace analysis using a gas chromatograph (HP6850, Agilent) equipped with a flame ionization detector and a megabore column (HP-PLOT Al^2^O^3^ S, 50m*0.53mm*0.15μm). The temperature for column chamber, inlet chamber and detector were 40 °C (isothermal), 150 °C and 220 °C, respectively. High purity hydrogen was used for carrier gas, at a flow rate of 2.6 mL.min^-1^. The split ratio of gas sample in inlet chamber was 25:1. The flow rate at the detector was 450mL.min^-1^ for air, 45mL.min^-1^ for hydrogen and 55 mL.min^-1^ for nitrogen. Methane volume and concentration in microcosms’ headspaces was calculated by comparing the areas of methane peaks obtained from the samples with a standard area, determined by the average of five injections of 99.95% pure methane (standard deviation < 1%). Clapeyron equation was used to calculate methane amounts in mmols, assuming temperature of 25 °C, atmospheric pressure (1 atm) and the volume corresponding to the total volume (milliliters) of methane consumed in each period.

## Results and Discussion

### Methanotrophs from sediment of Bertioga assessed by *pmo*A clone library

Methanotrophy is an important biological regulator of methane fluxes to the atmosphere. Aerobic processes were the first to be described. Anaerobic methane oxidation in hypoxic and microoxic natural and artificial environments [6, 34–36] were later detected, with the use of sulfate, nitrate, nitrite and metals as electron acceptors [7], showing that methanotrophs are able to occupy a number of diverse niches where methane is present. More recently, studies have been pointing out to the ability of aerobic methanotrophs to be active in anoxic environments competing with anaerobic groups [37].

A library of *pmo*A gene was carried out as a preliminary attempt to access the methanotrophic diversity in the mangrove sediment. The phylogenetic tree (Figure 1) reveals that 79% of the clones grouped with the gammaproteobaterial family *Methylococcaceae*, and the remaining 21% with the family *Methylocystaceae*, both already reported in anoxic environments [38]. Aerobic methanotrophic gammaproteobacteria are commonly detected in aquatic environments and in habitats rich in methane and at hypoxic or anaerobic conditions, indicating that there may be a niche overlap with anaerobic methanotrophs [37]. One example are species from the genus *Methylomonas*, capable of oxygen scavenging under hypoxic environments [37]. This genus was the most representative in the sediment used as inoculum, being affiliated to 21% of sequences. It has already been reported in the rhizosphere of Brazilian mangrove roots mesocosms [1]. Abundance of *Methylococcaceae* was also positively correlated with concentrations of hydrocarbons and negatively with dissolved oxygen in consequence of Deepwater Horizon disaster [39]. Given the fact that sequences of bacteria related to contaminated areas or able to hydrocarbon and MTBE degradation were detected in 16S rRNA libraries of the sediment (data not shown) and of SIP microcosms (Figures 4 and 5), it is possible that exposure of the microbiota to oil spills in the area of study may have contributed to the higher number of *Methylococcaceae* clones found.

**Fig. 1.**
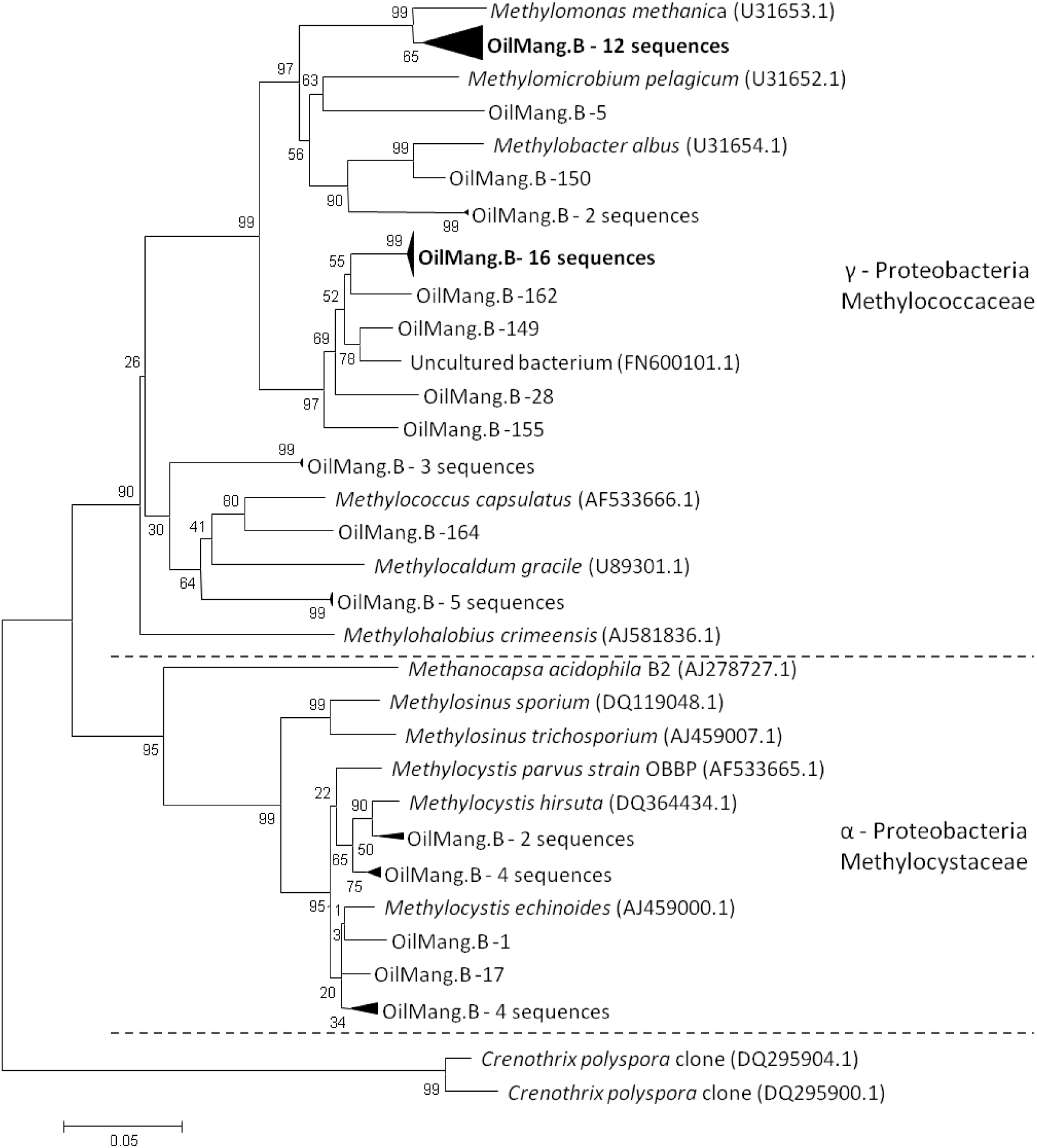
Phylogenetic neighbour-joining tree of *pmo*A gene sequences from clone library constructed from sediment of Bertioga’s mangrove, at oil impacted site (OilMang.B). Reference strains and clones sequences were taken from GenBank and are named by their accession numbers. Bootstrap values derived from 2000 replicates are shown and were obtained using a distance matrix program neighbour-joining method within MEGA 7.0. The bar represents 5% sequence divergence.

**Fig. 2.**
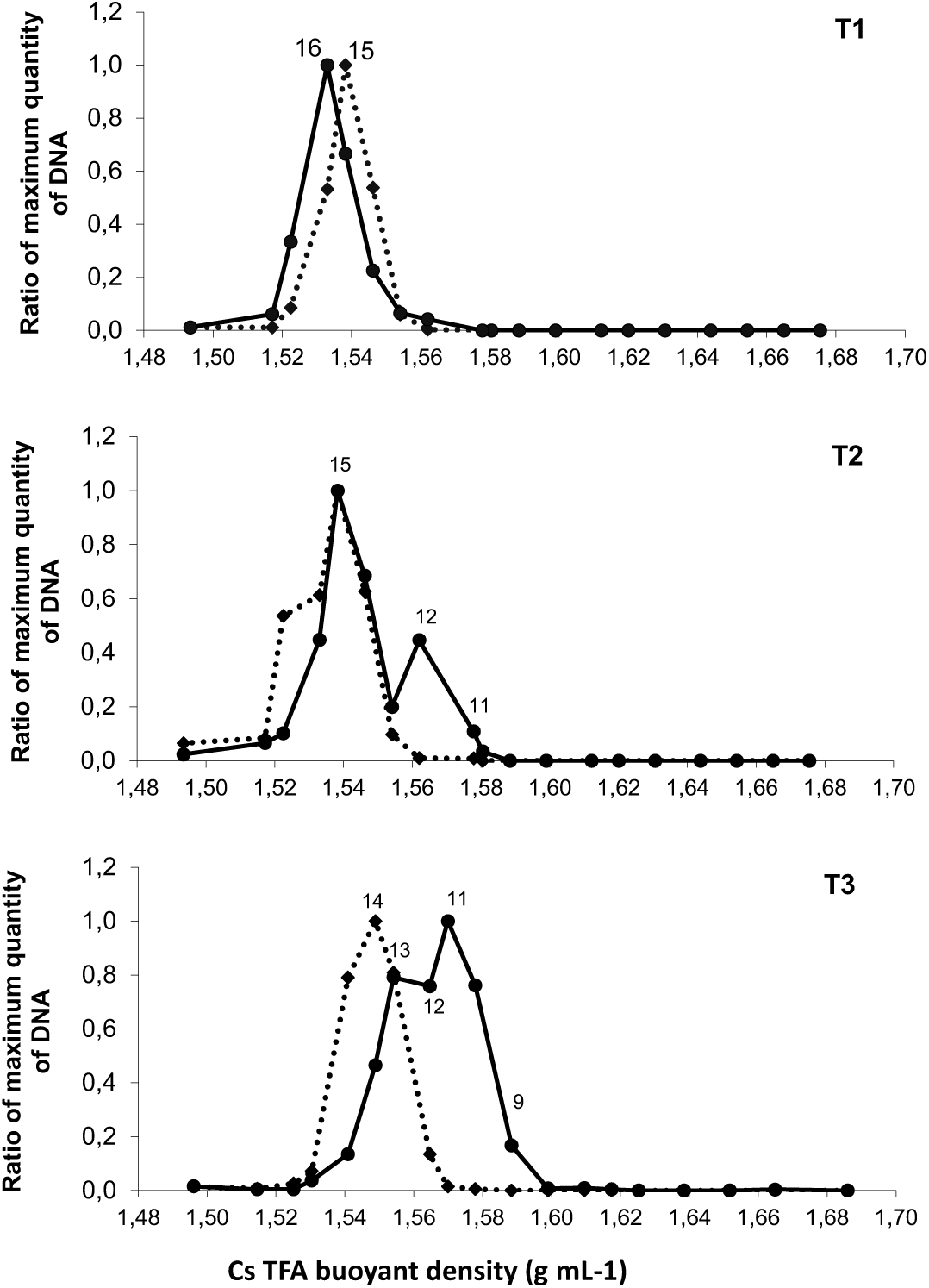
Quantitative distribution of DNA across the entire range of buoyant density (BD) of the DNA-SIP fractions from sediment microcosms incubated with 12CH4 (12C-DNA dotted line), and 13CH4 (13C-DNA solid line) for 2, 4 and 7 days of incubation (T1, T2 and T3, respectively). The numbers in the plot represent the fractions of prominence.

**Fig. 3.**
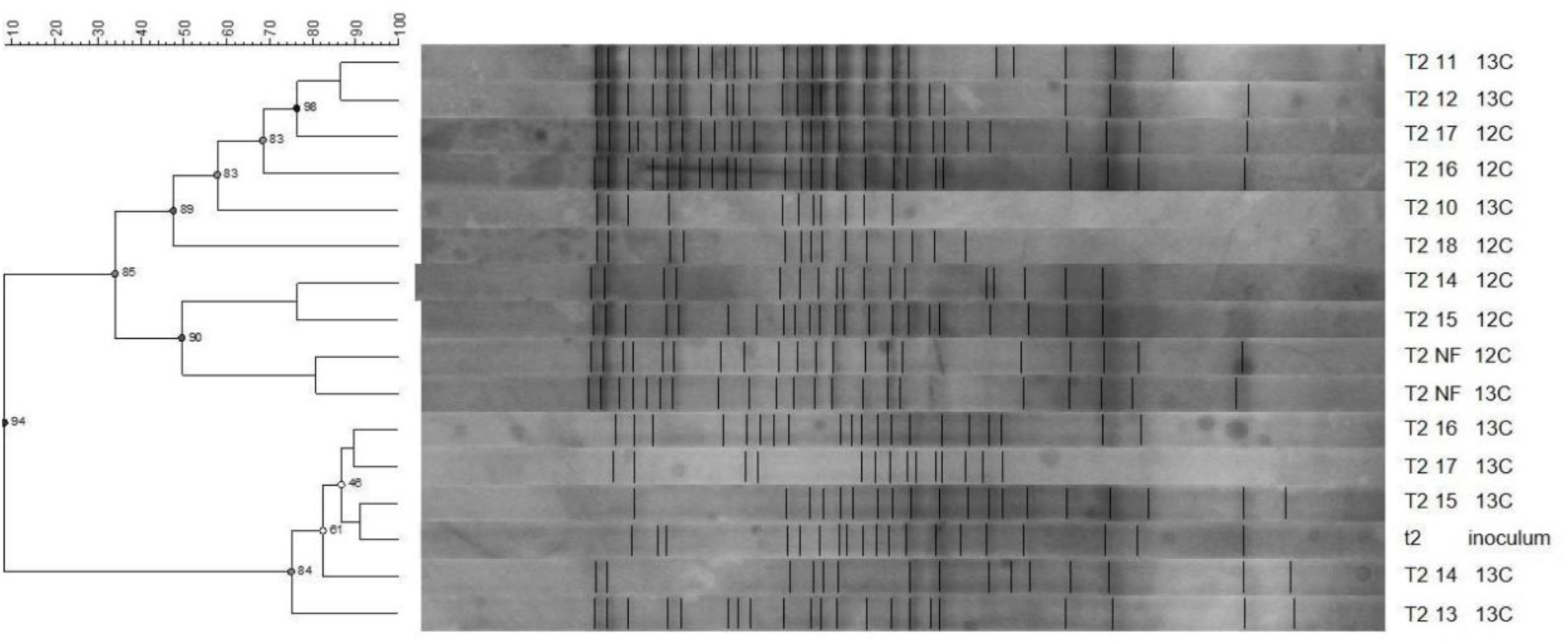
Cluster analysis of16S rDNA DGGE fingerprint of fractionated DNA recovered from 12CH4 and 13CH4 microcosms from mangrove samples (Bertioga, São Paulo State, Brazil), at the 4th day of incubation (T2). The denaturing gradient ranged from 45% to 65%. Cluster analysis was performed with the Bionumerics software (Applied Maths, NV), using UPGMA method and Pearson coefficient. Cophenetic correlation values are shown in the nodes. Labels in each lane indicate: incubation time (T2 = 4 days); number of recovered fractions after SIP-DNA experiment and isopycnic centrifugation; enrichment carbon source (12C = 12CH4 or 13C = 13CH4). t2, inoculum; NF, non-fractionated.

**Fig. 4.**
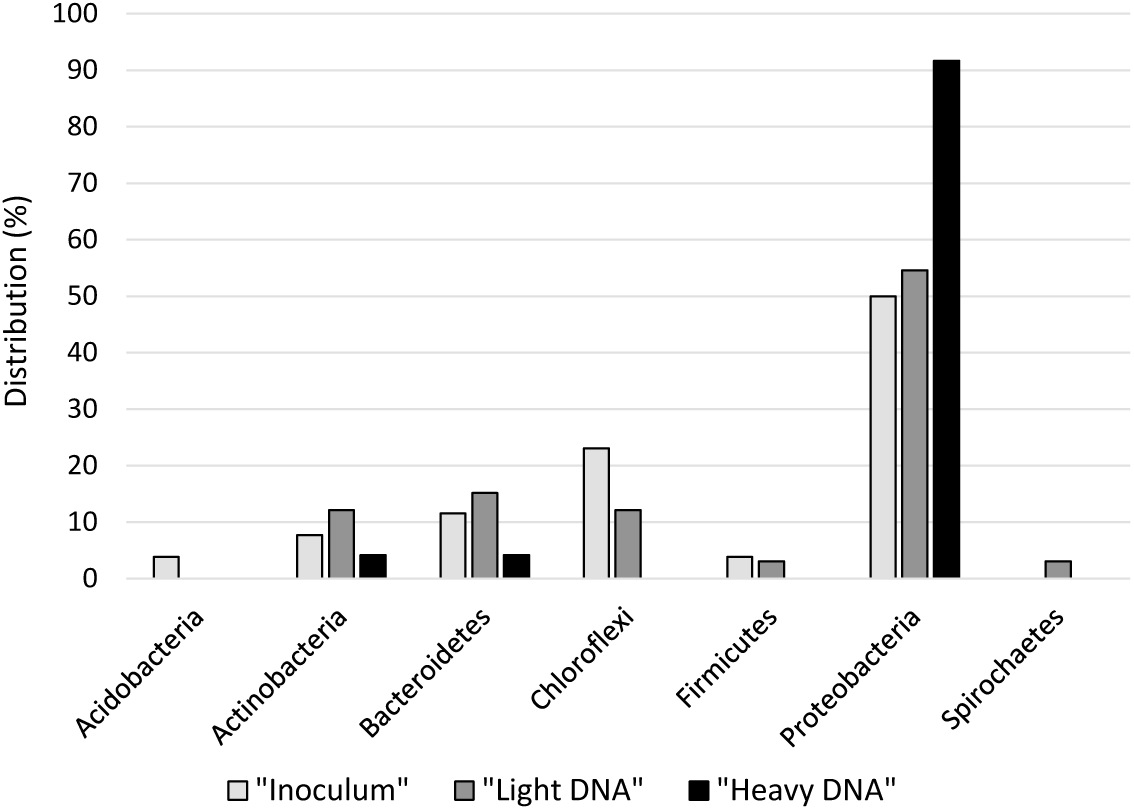
Phylum distribution detected by 16S RNA clone library from mangrove sediment (inoculum) and from light and heavy fractions derived from DNA-SIP experiment of microcosms enriched with methane (^13^CH_4_).

**Fig. 5.**
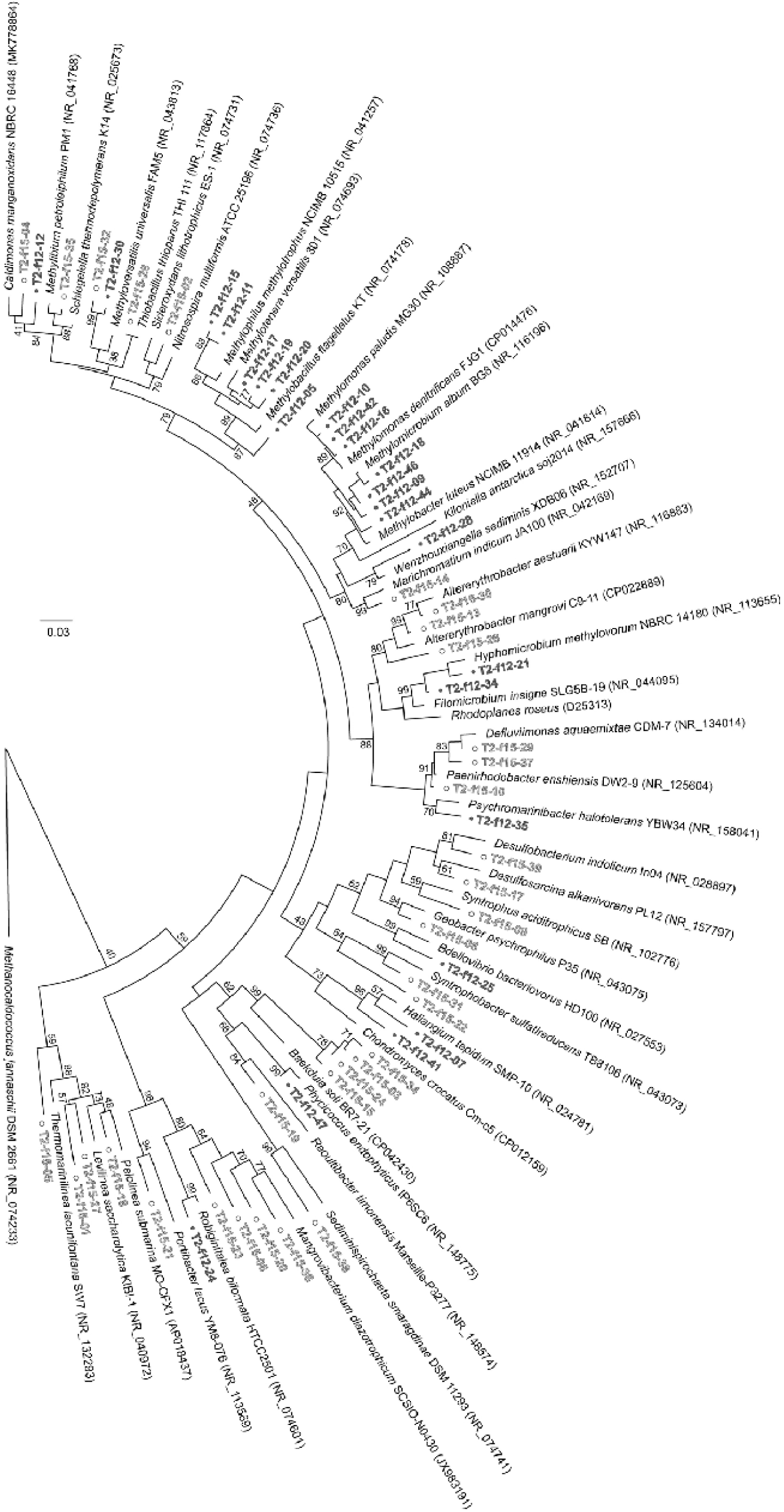
Phylogenetic tree of 16S rRNA gene, obtained from DNA of light (white circles) and heavy (black circles) fractions recovered from DNA-SIP experiment of mangrove sediment microcosms incubated with 13CH4. The maximum likelihood method with a 1,000 bootstrap analysis was used and representative reference sequences of the most closely related members were obtained from the Genbank and RDP.

There is also a comprehensive cluster whose nearest reference is an uncultured bacterium (FN600101.1) obtained from the rhizosphere of *Oryza sativa* in a rice field in Italy (unpublished). Indeed, uncultured methanotrophic groups have been observed in paddy fields in several parts of the world [40]. Moreover, 35% of sequences grouped with uncultured organisms demonstrating the lack of characterized isolates of the group.

In the set of sequences obtained there were not representatives of other groups of methanotrophic bacteria, such as *Crenothrix polyspora* and those included in the Phylum Verrucomicrobia. However, their presence in the sediments cannot be excluded, since i) library coverage may not have been complete and ii) these organisms have an unusual *pmo*A gene, which could only be accessed through the use of modified primers and/or less restrictive conditions of PCR [41, 42].

### DNA-SIP microcosms

The percentage of methane added to the microcosms was in agreement with other studies, which added 5 to 10% methane at each new supply [43–45]. In our study 0.7 mmol of methane was consumed in 4 days of incubation which is in agreement with slurries incubated with mineral medium [43] and this strategy is quite valuable to prevent the cross-feeding, very common especially after longer periods of incubation [44, 45].

Isopycnic density gradient centrifugation of DNA extracted from cultures at 2, 4 and 7 days of incubation (T1, T2 and T3, respectively) showed an unlabeled DNA peak at BD of 1.5383 and 1.5330 (fractions 15-16) whereas ^13^C-DNA occupied fractions ranging in BD from 1.5884 to 1.5620 (fractions 9-12) (Figure2). After 2 days of incubation, with consumption of 0.2 mmol of methane, no clear shift was obtained (Figure 2 T1). After 4 days of incubation (Figure 2 T2), DNA from fractions 11-12 was detected in the ^13^C-methane microcosm compared to the ^12^C-methane microcosm, showing incorporation of the ^13^C into the DNA of microbiota metabolically involved in methane oxidation. At this time, consumption of 0.7 mmol of methane was detected. After 7 days of incubation (Figure 2, T3) and 1.4 mmol methane consumption, DNA from the heaviest fractions were also detected in the ^13^C-methane microcosm compared to the ^12^C-methane microcosm, as well as DNA from light fractions (13-14). To minimize the influence of cross-feeding on the results, only DNA samples of the T2 were chosen for further analyses.

### Microbial community and phylogenetic analysis

PCR-based 16S rRNA gene DGGE analysis was carried out on the ^12^C- and ^13^C-enriched fractions of the microcosm after 4 days of incubation (Figure 3). PCR-based 16S rRNA gene DGGE analysis of inoculum as well as of the DNA not fractionated of the microcosm were also included (Figure 3). In relation to the inoculum (lane t2), it is possible to notice that after 4 days of incubation an enrichment of microorganisms involved in the metabolism of methane can be detected, evidenced by a cluster of NF C12 and NF C13 (DNA not fractionated of the flasks incubated with ^12^CH_4_ and ^13^CH_4_, respectively). Bacteria fingerprints of samples incubated with ^13^CH_4_ show that fractions 11-12 (T2 13C 11 and 12) formed a cluster with 88 % similarity. These fractions had an increase in DNA after ^13^CH_4_ addition, corresponding to the microorganisms that incorporated carbon from ^13^CH_4_. Interesting to note is that light fractions of the flask incubated with ^13^CH_4_ (T2 13C 13-17) representing microorganisms that did not incorporated carbon from ^13^CH_4_, formed a cluster with 78% similarity with the inoculum.

Clone libraries of the 16S rRNA gene were constructed from total DNA extracted from the mangrove sediment (inoculum), and from the fractions 12 (BD = 1.5620) and 15 (BD = 1.5383) of the ^13^C-methane amended microcosm, corresponding to “inoculum”, “heavy fraction”, and “light fraction” clone libraries, respectively. Analysis of the sequences against the RDPII database revealed that there was homology with 16S rRNA gene sequences from mangroves, marine sediments and xenobiotics contaminated sites. Phylum distribution of sequences in clone libraries from sediment and light fractions were similar, with predominance of Proteobacteria, Chloroflexi, Bacteroidetes and Actinobacteria (Figure 4). These phyla were also amongst the most abundant groups in Bertioga mangrove sediments metagenomes [16], corroborating the obtained results in this work of bacterial diversity in the area of study. On the other hand, as expected, incubation of sediment with methane resulted in the enrichment of the Proteobacteria phylum, as evidenced by the distribution of sequences in the heavy fraction library.

Sequences from light fractions (Figure 5, white circles) were widely distributed into different clades and mostly associated to chemoheterotrophic bacteria. Representative genera of sulfur (*Marichromatium*), iron (*Sideroxydans* and *Geobacter*) and manganese (*Caldimonas*) oxidizing bacteria as well as sulfate reducing bacteria (*Desulfobacterium*) were present. Detection of sequences associated to polyaromatic hydrocarbon degradation (*Alterythrobacter, Methylibium*) or to oil fields (*Defluvimonas*) may also reflect the adaptation of the microbiota to past oil spills in the area of study. The presence of anaerobic groups is probably associated to bacteria from the inoculum carried into the culture medium and still present after the short incubation period.

Sequences from the heavy fraction (Figure 5, black circles), on the other hand were grouped mainly in clades associated to methanotrophic (*Methylomonas, Methylomicrobium* and *Methylobacter*) and methylotrophic (*Methylotenera, Methyloversatilis, Methylophilus* and *Hyphomicrobium*) groups, reflecting the associations of bacterial groups related to the methane cycle in the Bertioga mangrove sediment and in agreement with a possible methane derived microbial food chain [37]. The recovery of methylotrophic bacteria sequences in the heavy DNA fractions, in particular β-Proteobacteria, have also occurred in several studies involving the DNA-SIP technique [44–47]. Recently, cooperation between *Methylobacter* (*Methylococcaceae*) and *Methylotenera versatili* (*Methylophilaceae*) on methane oxidation was evidenced in lake Washington sediment fed with ^13^C-methane and incubated under aerobic conditions and with nitrate (10mM) as electron acceptor [48]. A succession of methanotrophic and methylotrophic metabolisms coupled to denitrification was also described in a membrane biofilm reactor treating perchlorate and fed with methane as sole electron donor and carbon source [49]. Initially performing anaerobic methanotrophy coupled to denitrification, the reactor biofilm was gradually dominated by *Methylocysytis* and at final stages, by *Methylomonas* cells. Under low dissolved oxygen tensions, these genera coupled methane oxidation to nitrate reduction (intraerobic methanotrophy), generating nitrite or nitrous oxide. The organic compounds derived from methanotrophy, such as methanol, were oxidized by the methylotrophic denitrifying genera *Methylophilus* and *Methyloversatilis* coupled to nitrate reduction. The authors also hypothesized that anaerobic methanotrophic microorganisms could benefit from intermediates produced by aerobic methanotrophs and both metabolisms occurred simultaneously under this methane rich anoxic environment. Similar associations may be occurring at the sediments of Bertioga mangrove. In our study, *Methylomonas, Methylobacter and Methylotenera* genera were dominant in the heavy fraction of DNA-SIP microcosms. NMS medium is rich in nitrate and, although incubations were carried out under aerobiosis, the concomitant presence of these groups in the microcosms associated to ^13^CH_4_ consumption may indicate that under the anoxic environment of the mangrove sediment, they could shift to intraerobic methane oxidation. Another example of methylotrophic denitrifying bacterium commonly associated to methanotrophic bacteria in the heavy fraction sequences was *Hyphomicrobium methylovorum*. The fact that the genus *Methylocystis*, retrieved from sediment in pmoA clone libraries, was not detected in DNA-SIP microcosms may be related to the influence of culture conditions, probably favouring *Methylomonas* over *Methylocystis* cells.

Other clones from the heavy fraction clone library occurred at low frequency (Fig. 5). These clones might be representatives of: i) bacteria that incorporated heavy carbon through the use of secondary marked substrates such as methanol and formaldehyde or ii) sequences amplified from light DNA traces [45, 47].

## Conclusion

Our work showed that aerobic bacterial community associated to methane consumption in Bertioga sediment is diverse and that they may have an important role in reducing methane emissions under aerobic or anoxic conditions. Both alpha and betaproteobacteria methanotrophs commonly associated to aquatic environments were represented in *pmo*A clone libraries, with predominance of the Methylococcaceae family, in special *Methylomonas* and *Methylobacter* genera. Methylocystaceae family was also represented by *Methylocystis* sequences and a clade of sequences related to an uncultured clone sequence also indicate that other methanotrophic groups are yet to be identified.

DNA-SIP technique, followed by analysus of 16S rRNA gene, was a suitable tool to detect active microorganisms related to methane oxidation, using microcosms with NMS medium containing nitrate and amended with methane as sole source of carbon and energy. Sequences from light fractions were broadly distributed into clades of chemoheterotrophic bacteria, including groups associated to hydrocarbon degradation or to oil fields, as well as genera related to sulfur, iron and manganese oxidizing and sulfate reducing metabolisms. Microbial groups involved in ^13^CH_4_ consumption were mostly methanotrophic (*Methylomonas, Methylomicrobium* and *Methylobacter*) and methylotrophic (*Methylotenera, Methyloversatilis, Methylophilus*, and *Hyphomicrobium*) bacteria. Their co-occurrence in DNA-SIP microcosms suggests that a microbial methane food chain may be established in Bertioga sediments and it is possibly able to shift from aerobic methanotrophy to methane oxidation coupled to nitrate reduction when oxygen concentrations are low.

## Compliance with Ethical Standards

The authors declared that they have no conflicts of interest to this work. We declare that we do not have any commercial or associative interest that represents a conflict of interest in connection with the work submitted.

## Acknowledgments

We thank the research team from USP and EMBRAPA, Rosa Gamba and Dr. Armando C. F. Dias for their scientific, logistic and sampling support.

